# V3 tip determinants of susceptibility to inhibition by CD4-mimetic compounds in natural clade A human immunodeficiency virus (HIV-1) envelope glycoproteins

**DOI:** 10.1101/2023.07.31.551312

**Authors:** Saumya Anang, Shijian Zhang, Christopher Fritschi, Ta-Jung Chiu, Amos B. Smith, Navid Madani, Joseph Sodroski

**Affiliations:** Department of Cancer Immunology and Virology, Dana-Farber Cancer Institute, Boston, Massachusetts, USA; Department of Microbiology, Harvard Medical School, Boston, Massachusetts, USA; Department of Chemistry, University of Pennsylvania, Philadelphia, Pennsylvania, USA

**Keywords:** virus, envelope, entry inhibitor, Env, CD4-mimetic compound, resistance, gp120, variable region, V3 loop, strain variation

## Abstract

CD4-mimetic compounds (CD4mcs) bind the human immunodeficiency virus (HIV-1) gp120 exterior envelope glycoprotein (Env) and compete for binding to CD4, the host receptor. CD4mcs prematurely trigger conformational changes in Env similar to those induced by CD4, leading to transient activation of infectivity followed by irreversible virus inactivation. Natural HIV-1 variants exhibit a wide range of susceptibilities to CD4mc inhibition, only a small fraction of which can be explained by variation in the gp120 Phe-43 cavity/vestibule where CD4mcs bind. Here, we study Envs from the resistant HIV-1_BG505_ and the more sensitive HIV-1_191955_A4_ clade A strains. The major determinant of the relative sensitivity of the HIV-1_191955_A4_ Env to CD4mcs mapped to a single residue change (F317Y) in the tip of the gp120 V3 variable loop. In the Envs of several HIV-1 strains, replacement of the more prevalent Phe 317 with a tyrosine residue increased virus sensitivity to multiple CD4mcs. Tryptophan substitutions at residues 317 and 316 resulted in increases and decreases, respectively, in sensitivity to CD4mcs. Some of the gp120 V3 changes increased virus sensitivity to inactivation by both CD4mc and cold exposure, phenotypes indicative of increased Env triggerability. Infection of CD4-negative cells expressing the CCR5 coreceptor by these Env variants was triggered more efficiently by CD4mcs. For the panel of studied HIV-1 Envs, resistance to the CD4mcs was associated with decreased ability to support virus entry. These studies illustrate how variation in gp120 outside the CD4mc binding site can influence the sensitivity of natural HIV-1 strains to inhibition by these compounds.

**IMPORTANCE:** CD4-mimetic compounds (CD4mcs) are small-molecule inhibitors of human immunodeficiency virus (HIV-1) entry into host cells. CD4mcs target a pocket on the viral envelope glycoprotein (Env) spike that is used for binding to the receptor, CD4, and is highly conserved among HIV-1 strains. Nonetheless, naturally occurring HIV-1 strains exhibit a wide range of sensitivities to CD4mcs. Our study identifies changes distant from the binding pocket that can influence the susceptibility of natural HIV-1 strains to the antiviral effects of multiple CD4mcs. We relate the antiviral potency of the CD4mc against this panel of HIV-1 variants to the ability of the CD4mc to activate entry-related changes in Env conformation prematurely. These findings will guide efforts to improve the potency and breadth of CD4mcs against natural HIV-1 variants.

## INTRODUCTION

The binding of the human immunodeficiency virus (HIV-1) envelope glycoprotein (Env) to host cell receptors, CD4 and CCR5/CXCR4, triggers virus entry into the cell (1–11). The Env trimer consists of three gp120 exterior Envs and three gp41 transmembrane Envs. Prior to receptor engagement, the HIV-1 Env trimer on virions mainly exists in a pretriggered “closed” conformation (State 1), but also samples more “open” conformations (States 2 and 3) (12–15). CD4 binding drives Env from State 1 to State 2 and then into State 3, the prehairpin intermediate (12–18). In the prehairpin intermediate, the heptad repeat (HR1) region of gp41 forms an exposed coiled coil (16–19). Binding of the State 3 Env to the CCR5 or CXCR4 coreceptor is thought to induce the formation of a highly stable gp41 six-helix bundle, which promotes the fusion of the viral and cell membranes (20–25).

The closed nature of the State-1 conformation, significant strain variability and heavy glycosylation of the HIV-1 Env trimer contribute to the avoidance of potentially neutralizing antibodies (26–29). Amidst conformational and sequence variation, HIV-1 must conserve the gp120 binding sites for its receptors. The binding site for CD4 on gp120 consists of a conserved surface that is conformationally altered by CD4 binding. CD4 binding creates an internal pocket in gp120 called the Phe-43 cavity that is bounded by highly conserved residues from gp120 and a single phenylalanine residue (Phe 43) from CD4 (30). The ∼150-Å^3^ Phe-43 cavity and the surrounding “vestibule” on the gp120 surface comprise the binding sites for two classes of small-molecule HIV-1 entry inhibitors, the CD4-mimetic compounds (CD4mcs) and conformational blockers like BMS-806 and BMS-529 (temsavir) (30–40).

CD4-mimetic compounds (CD4mcs) disrupt HIV-1 entry by binding to gp120 in the Phe-43 cavity, directly competing with CD4 but also prematurely triggering Env (31, 40–42). CD4mcs drive the State-1 Env trimer into downstream conformations (States 2 and 3) that, in proximity to a coreceptor-expressing target cell, can mediate HIV-1 infection (42). These CD4mc-induced Env intermediates are short-lived and, in the absence of a coreceptor-expressing target cell, irreversibly decay into inactive, dead-end conformations (41, 42). At CD4mc concentrations that do not completely inhibit HIV-1 infection, the induction of more open Env conformations sensitizes HIV-1 viruses to neutralization and HIV-1-infected cells to antibody-dependent cellular cytotoxicity (ADCC) by otherwise ineffectual antibodies (43–52).

Early CD4mcs discovered using a gp120-CD4 screen exhibited weak antiviral potency against a limited range of HIV-1 isolates (31, 40). Iterative cycles of design, guided by CD4mc-gp120 structures and empirical testing, led to the development of analogues with improved potency (36–38, 53–56). Progressive increases in CD4mc potency have been accompanied by an increase in the breadth of activity against a wider range of HIV-1 strains (37, 38, 57). BNM-III-170, a well-studied CD4mc analogue with an indane scaffold, inhibits approximately 70% of a global panel of multi-clade HIV-1 variants (37). Recently developed CD4mcs based on an indoline scaffold exhibit 10-20-fold increases in anti-HIV-1 potency compared with BNM-III-170 (57). With the exception of CRF01_AE recombinant HIV-1 (see below), indoline CD4mcs inhibit the entry of every HIV-1 strain tested (57).

Despite significant improvements in the potency and coverage of the lead indoline CD4mcs, primary HIV-1 strains exhibit a 1000-fold range of sensitivities to their antiviral effect (57). The rank order of sensitivities of lentivirus vectors pseudotyped by diverse HIV-1 Envs is highly correlated among different CD4mcs, suggesting that an intrinsic property of Env determines CD4mc susceptibility (57). With respect to HIV-1 phylogeny, only the CRF01_AE recombinants, in which the imidazole ring of His 375 occupies the Phe-43 cavity, are resistant to the CD4mcs (58–60). In the other phylogenetic clades, HIV-1 strains exhibit the entire range of sensitivities to the CD4mc (57). The susceptibility of most primary HIV-1 strains to CD4mcs is not obviously explained by local variation in the known binding site of these compounds (36–38, 57, 59–61). For example, the Phe-43 cavity of 96% of these HIV-1 strains is bounded by Ser or Thr 375 residues that are compatible with efficient CD4mc binding (58–61).

Pathways to CD4mc resistance preferred by HIV-1 in the absence of immune selection have been studied by passaging HIV-1 in the presence of BNM-III-170 (62). In addition to two changes near the gp120 Phe-43 cavity, a third change in the gp120 inner domain outside the BNM-III-170 binding site contributed to resistance. Studies with closely matched Env mutants have provided insight into one mechanism whereby Env changes distant from the CD4mc binding site can influence virus sensitivity to these compounds (26, 63–66). To bind and inhibit HIV-1, CD4mcs must induce transitions from State 1 to downstream conformations (26, 31, 35–37, 41, 42, 46, 47, 53–55, 67). Viruses with Envs that have more stable State-1 conformations and therefore are less prone to make transitions from State 1 exhibit greater resistance to CD4mcs (26, 63–66). Thus, Env “triggerability” or intrinsic reactivity, a property that is inversely related to the height of the activation barrier separating State 1 and State 2 (13, 26, 27), can significantly affect the susceptibility of HIV-1 Env mutants to inhibition by CD4mcs.

Little is known about the basis for the 1000-fold range of sensitivities of natural non-CRF01_AE HIV-1 strains to the lead indane and indoline CD4mcs. CD4mcs make backbone contacts or interact with the side chains of highly conserved gp120 residues (37, 55, 57). Thus, an alternative explanation of primary HIV-1 susceptibility involving differences in Env triggerability, which need not be specified by changes near the CD4mc binding site, is appealing. Here, we investigate the basis for the different sensitivity of two clade A primary viruses, HIV-1_191955_A4_ and HIV-1_BG505_, to BNM-III-170 and potent lead indoline CD4mcs. We map the Env determinant of this difference in CD4mc sensitivity to a single amino acid residue (Tyr/Phe 317) in the tip of the gp120 V3 loop. We explore the effect of changes in this and adjacent V3 residues on the sensitivity of viruses to CD4mcs and cold exposure, two phenotypes known to be influenced by alterations in Env triggerability (63–66, 68). We evaluate the impact of Env resistance to CD4mcs on the ability to mediate infection of cells expressing CD4 and CCR5. We also examine the ability of the Env variants to be activated by CD4mcs to mediate virus infection of CD4-negative, CCR5-expressing cells. These results provide insights into mechanisms whereby Env changes outside the CD4mc binding site can influence the sensitivity of natural HIV-1 strains to inhibition by this class of entry inhibitors.

## RESULTS

### Determinants of the different sensitivities of the 191955_A4 and BG505 Envs to CD4mcs

HIV-1_191955_A4_, hereafter referred to as HIV-1_A4_, and HIV-1_BG505_ are primary clade A viruses (69–71) that exhibit differences in sensitivity to several CD4mcs (Fig. 1A). Compared with HIV-1_BG505_, HIV-1_A4_ is relatively sensitive to the indane CD4mc BNM-III-170 (37) and to the indoline CD4mcs, CJF-III-192 and CJF-III-288 (57). As expected (57), the indoline CD4mcs inhibited both HIV-1_A4_ and HIV-1_BG505_ more potently than BNM-III-170. As there are more than 120 amino acid residue differences between the A4 and BG505 Env ectodomains, we first attempted to localize the determinants of CD4mc sensitivity to the gp120 or gp41 subunit. The gp120_B_gp41_A_ chimera, which comprises the BG505 gp120 and A4 gp41, exhibited a level of resistance to the CD4mcs at least as great as that of the BG505 parent virus (Fig. 1A). Conversely, the gp120_A_gp41_B_ chimera was as sensitive to inhibition by the CD4mcs as the A4 parent virus. These results indicate that main determinants of the differences in CD4mc susceptibility between the A4 and BG505 viruses are located in the gp120 subunit.

**Fig 1:**
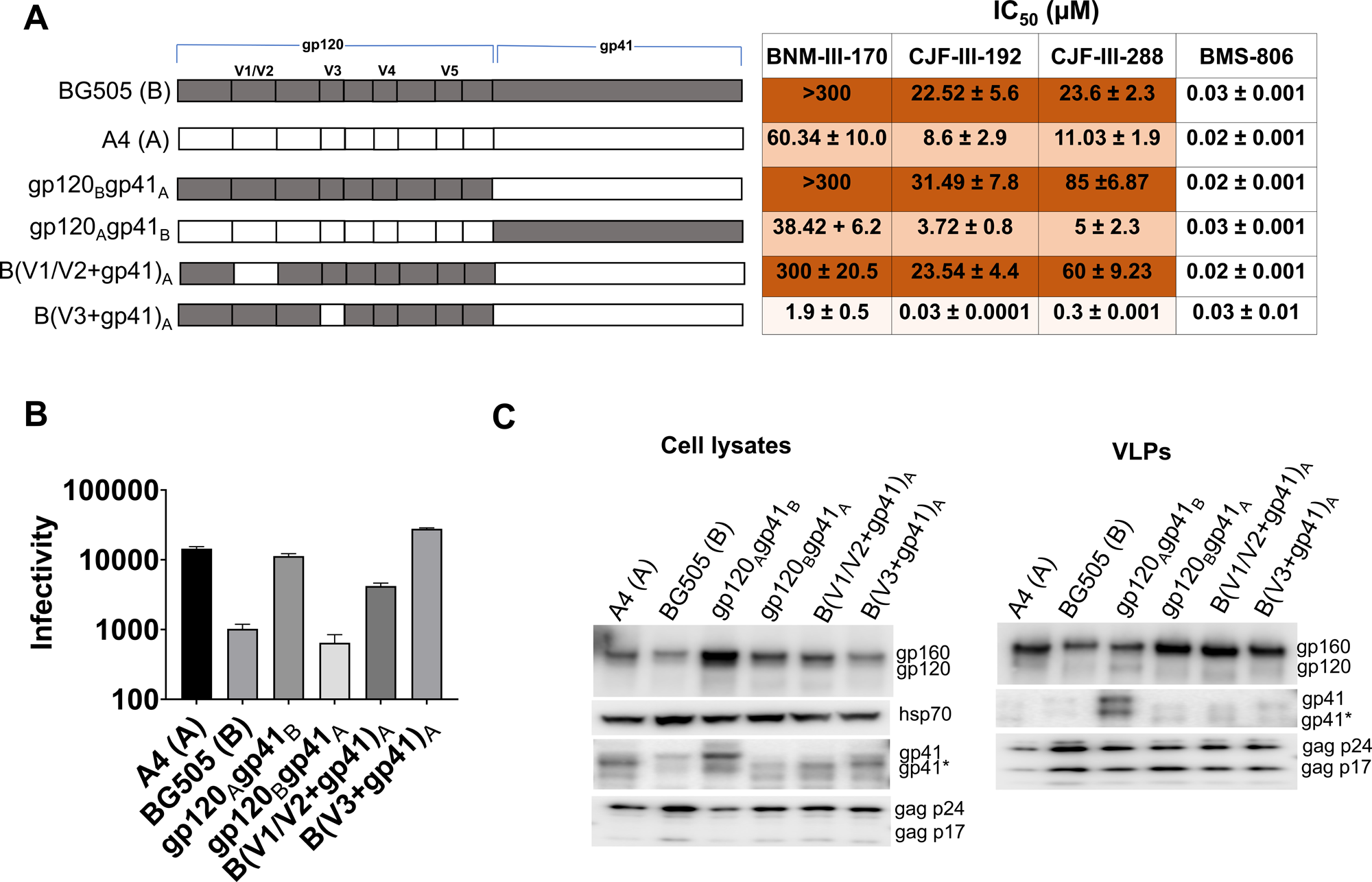
Determinants of the different sensitivities of the 191955_A4 and BG505 Envs to CD4mcs. (A) A schematic representation is shown of the gp120 and gp41 subunits of the 191955_A4 (designated A4) and BG505 Envs and chimeras. The gp120 V1/V2, V3, V4, and V5 variable regions are designated. HEK 293T cells were transfected with plasmids encoding the indicated Envs, HIV-1 packaging proteins and a luciferase-expressing HIV-1 vector. Pseudoviruses were incubated with the different concentrations of the inhibitors for 1 hour at 37°C and then added to Cf2Th-CD4/CCR5 cells. After 2 days of culture, the cells were lysed, the luciferase activity was measured and the IC_50_ was calculated. The IC_50_ values for inhibition of viruses pseudotyped with the indicated Envs by BNM-III-170, CJF-III-192, CJF-III-288 and BMS-806 are reported as means and standard deviations derived from at least three independent experiments. The intensity of shading is proportionate to the level of CD4mc resistance. (B) The infectivity of viruses with the indicated Envs was measured on Cf2Th-CD4/CCR5 cells. Recombinant luciferase-expressing viruses pseudotyped with the indicated Envs were produced as described above. Forty-eight hours later, cell supernatants were cleared and added to Cf2Th-CD4/CCR5 cells. Two days later, the cells were lysed and the luciferase activity was measured. (C) The expression level, processing and virion incorporation of the A4 and BG505 Envs and indicated chimeras are shown. HEK 293T cells were transfected with the packaging plasmids and constructs expressing the Env variants. Cell lysates and virus-like particles (VLPs) were harvested 48 hours later and analyzed by Western blotting for the indicated proteins. A minor form of gp41 resulting from cleavage of the cytoplasmic tail is designated by an asterisk. The results shown are representative of those obtained in three or more independent experiments. However, the relatively high level of gp41 in the VLPs containing the gp120_A_gp41_B_ chimera was not reproduced in other experiments.

The testing of additional A4-BG505 chimeras involving different regions of gp120 allowed us to narrow the number of potential candidates for the CD4mc susceptibility determinants further. The results with one such chimeric Env, B(V1/V2+gp41)_A_, indicated that the V1/V2 region of BG505 gp120 does not determine the resistant phenotype (Fig. 1A). By contrast, the results with the B(V3+gp41)_A_ chimera clearly indicate that the BG505 gp120 V3 region is required for the relative resistance of the BG505 Env to the CD4mcs (Fig. 1A). The parental A4 and BG505 viruses and the viruses with chimeric Envs were comparably inhibited by the conformational blocker, BMS-806 (Fig. 1A). Thus, even though CD4mcs and BMS-806 bind to similar gp120 regions (30–40), sensitivity to these entry inhibitors differs among these Env variants.

The A4-BG505 chimeric Envs exhibited different levels of efficiency with which they supported virus infection of CD4^+^CCR5^+^ cells (Fig 1B). These differences were not related to variation in the level of Env expression or incorporation into virus particles (Fig. 1C). The relationship between CD4mc resistance and infectivity will be explored for these and other Env variants below.

### The gp120 V3 loop as a determinant of CD4mc susceptibility

The above results indicate that BG505-specific sequences in the gp120 V3 region are necessary for the relative resistance of HIV-1_BG505_ to the CD4mcs. We therefore tested whether particular BG505 V3 amino acid residues are essential for maintaining the resistant phenotype. Comparison of the A4 and BG505 gp120 V3 sequences identified six amino acid differences (Fig. 2A). Our attention was drawn to the four differences at residues 307/308 and 317/318, which are symmetrically positioned at the tip of the V3 loop (72). While polymorphisms in three of these V3 residues are common in HIV-1 strains, Tyr 317 in the A4 Env is unusual (59, 60). Approximately 79% of HIV-1 strains have a phenylalanine residue at this position; in the remaining strains, Leu and Trp are common substitutions, Met is less common and Tyr is only occasionally seen. We evaluated the effect of introducing the F317Y change in the BG505 Env. Viruses with the B(307+317)_A_ and B(317)_A_ Envs were more sensitive to the CD4mcs than the BG505 virus, with sensitivities similar to or even greater than those of the A4 virus (Fig. 2A). Thus, the Tyr 317 substitution in V3 is sufficient to explain the increased CD4mc sensitivity of the A4 virus relative to that of the BG505 virus.

**Fig 2:**
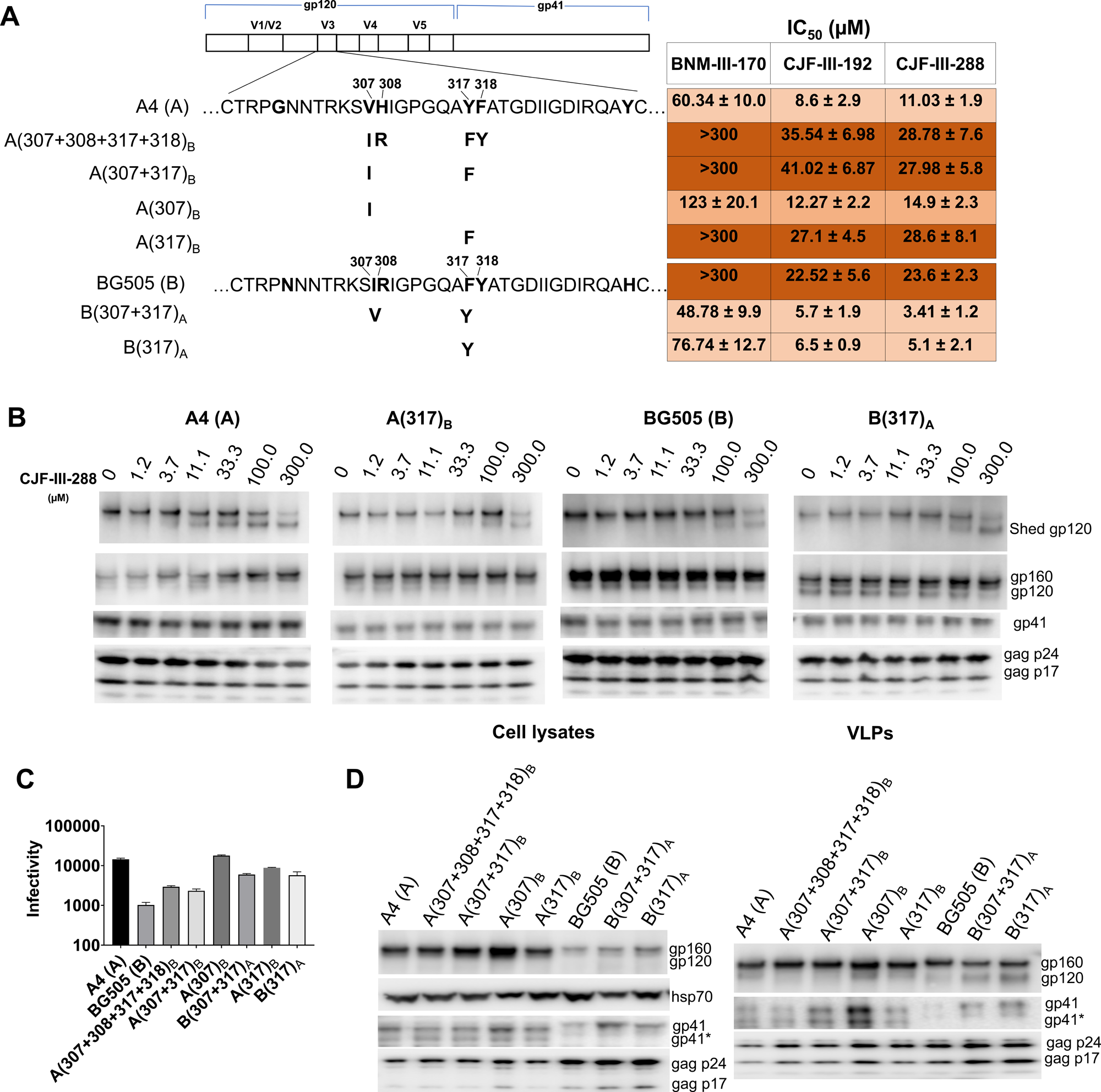
Analysis of the gp120 V3 loop as a determinant of CD4mc susceptibility. (A) The sequences of the A4 and BG505 gp120 V3 loops are aligned beneath the schematic representation of the HIV-1 Env. Amino acid residues that differ between the A4 and BG505 Envs are in bold. The amino acid residues changed in each Env variant are shown in bold beneath the parental Env. Standard HIV-1 Env amino acid numbering is used here and throughout the manuscript (89). The IC_50_ values for inhibition of infection by the CD4mcs were calculated as described in the Fig. 1A legend and are reported as means and standard deviations derived from at least three independent experiments. The intensity of shading is proportionate to the level of CD4mc resistance. (B) CJF-III-288-induced shedding of gp120 from virions with the indicated Envs was measured. Virions were produced transiently from HEK 293T cells transfected with pNL4-3 proviral constructs containing the A4, BG505, A(317)_B_ and B(317)_A_ *env* genes. Viruses were harvested, clarified by low-speed centrifugation, filtered (0.45-μm) and pelleted at 14,000 × *g* for 1.5 hours at 4°C. The virus pellet was resuspended in 1X PBS and incubated with the indicated concentrations of CJF-III-288 for 2.5 hours at 37°C. The viruses were then pelleted; the virus pellet was lysed and the supernatant containing shed gp120 was incubated with Galanthus nivalis lectin (GNL) beads. The viral lysates and the proteins captured on the GNL beads were Western blotted to detect the indicated HIV-1 proteins. (C) The infectivity of viruses with the indicated Envs is shown. The infectivity was measured on Cf2Th-CD4/CCR5 cells as described in the legend to Fig. 1B. (D) The expression level, processing and virion incorporation of the indicated Envs were evaluated as described in the Fig. 1C legend.

We evaluated whether the BG505 V3 sequences were sufficient to render the A4 virus resistant to the CD4mcs. The A(307+308+317+318)_B_, A(307+317)_B_ and A(317)_B_ viruses were as resistant to the CD4mcs as the BG505 virus (Fig. 2A). The level of gp120 shedding from the virus particles induced by CJF-III-288 roughly corresponded to the sensitivity of the viruses to inhibition by the CD4mcs (Fig. 2B). The A(307)_B_ virus exhibited an intermediate level of resistance to the CD4mcs. The expression and ability to support infection of these Envs are shown in Fig. 2C and D. Along with the above results, these data suggest that variation in the gp120 V3 tip is both necessary and sufficient to account for the different susceptibilities of the A4 and BG505 viruses to CD4mcs.

### Effect of V3 changes on CD4mc sensitivity in other HIV-1 strains

We asked whether the alteration of Phe 317 to the less-common tyrosine residue would alter CD4mc sensitivity in other HIV-1 strains. We introduced the F317Y change into the Envs of HIV-1_191084_, another clade A virus, and HIV-1_CRF02_AG253.11_, a Tier-3 clade AG recombinant virus (69, 73–75). In both HIV-1 strains, the F317Y change resulted in increased sensitivity to the CD4mcs (Fig. 3A). Thus, the effect of substitution of a tyrosine residue at Phe 317 on CD4mc sensitivity applies to multiple HIV-1 strains. The expression and ability to support infection of these Envs are shown in Fig. 3B and C. We note that the wild-type AG253.11 Env was less efficient in supporting virus infection than the other Envs studied, despite a comparable level of Env expression and incorporation into virus particles.

**Fig 3:**
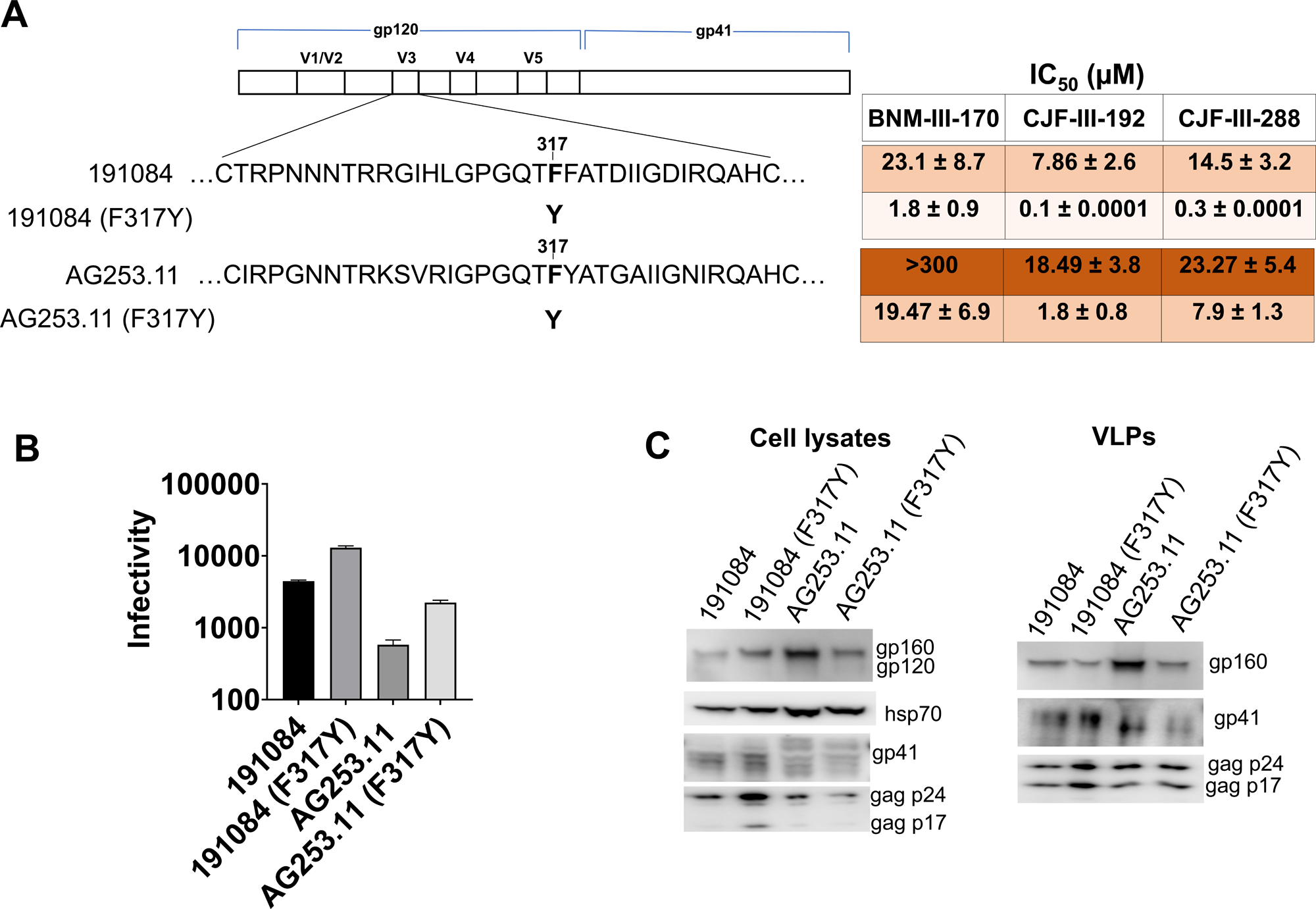
Effect of changes in V3 residue 317 on CD4mc sensitivity in other HIV-1 strains. (A) Sequences of the V3 loops of the 191084 and AG253.11 Envs and F317Y mutants are shown beneath the schematic of the HIV-1 Env. The IC_50_ values for inhibition of infection by the CD4mcs were calculated as described in the Fig. 1A legend and are reported as means and standard deviations derived from at least three independent experiments. The intensity of shading is proportionate to the level of CD4mc resistance. (B) The infectivity of viruses with the indicated Envs is shown. The infectivity was measured on Cf2Th-CD4/CCR5 cells as described in the legend to Fig 1B. (C) The expression level, processing and virion incorporation of the indicated Envs were evaluated as described in the legend to Fig. 1C.

Hydrophobic aromatic or aliphatic amino acids are found at residue 317 in most HIV-1 strains (59, 60). We asked if a hydrophobic Trp residue at 317 or the adjacent 316 position would influence HIV-1 susceptibility to inhibition by CD4mcs. The A316W change has been used to decrease the exposure of the V3 loop on soluble gp140 SOSIP.664 trimers (76, 77). Although the BG505 (F317W) virus was inhibited by CD4mcs comparably to the wild-type BG505 virus, the Y317W change in the A4 Env and the F317W change in the AD8 Env rendered the viruses more sensitive to the CD4mcs (Fig. 4A). By contrast, viruses with the A316W change were slightly or moderately more resistant to CD4mcs. These results indicate that changes in the tip of the gp120 V3 loop can affect the susceptibility of viruses from multiple HIV-1 strains to inhibition by CD4mcs. The expression and ability to support infection of these Envs are shown in Fig. 4B and C.

**Fig 4:**
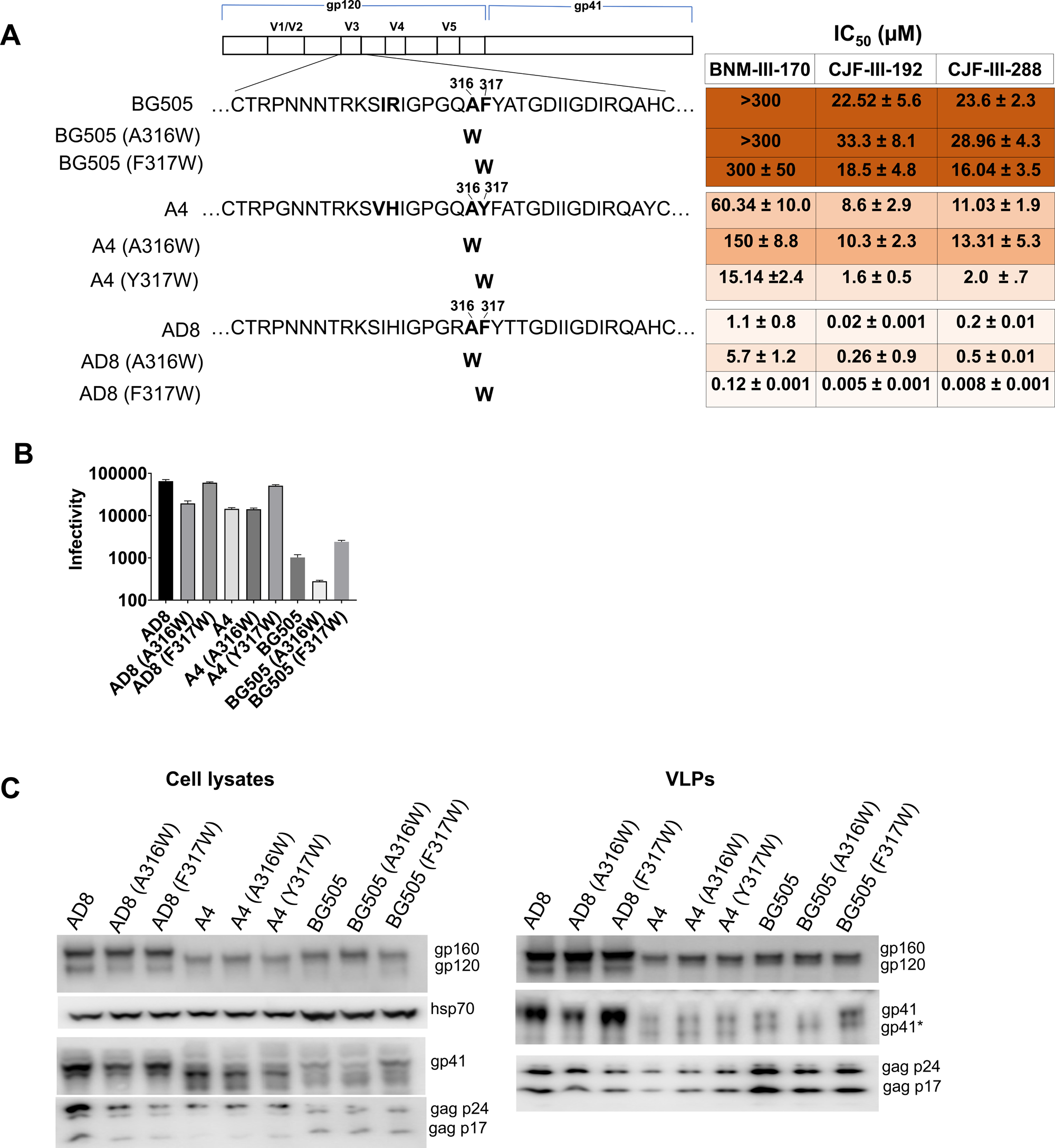
Effect of hydrophobic changes in the gp120 V3 loop. (A) Sequences of the gp120 V3 region of the BG505, A4 and AD8 Envs are shown, along with the hydrophobic Trp substitutions made in the V3 loop mutants, beneath the HIV-1 Env schematic. The IC_50_ values for inhibition of infection by the CD4mcs were calculated as described in the Fig. 1A legend and are reported as means and standard deviations derived from at least three independent experiments. The intensity of shading is proportionate to the level of CD4mc resistance. (B) The infectivity of viruses with the indicated Envs is shown. The infectivity was measured on Cf2Th-CD4/CCR5 cells as described in the legend to Fig. 1B. (C) The expression level, processing and virion incorporation of the indicated Envs were evaluted as described in the legend to Fig. 1C.

### CD4 activation of mutant virus infection

CD4mcs induce short-lived, activated Env intermediates that irreversibly decay into dead-end conformations (31, 41, 42). To investigate whether V3 loop changes affect this process, we examined the ability of the viruses with wild-type and mutant Envs to infect CD4-negative, CCR5-expressing cells in the presence of increasing concentrations of the CD4mcs. The activating effects of BNM-III-170 and CJF-III-288 were similar (Fig. 5A and B). Compared with the wild-type BG505, AD8, 191084 and AG253.11 viruses, the viruses with Tyr or Trp substitutions at residue 317 were generally activated more efficiently by the CD4mcs. Conversely, the Y317F change in the A(317)_B_ mutant decreased CD4mc activation relative to that of the wild-type A4 virus. Compared with the wild-type AD8 virus, viruses with the A316W Env were less efficiently activated by CD4mcs.

**Fig 5:**
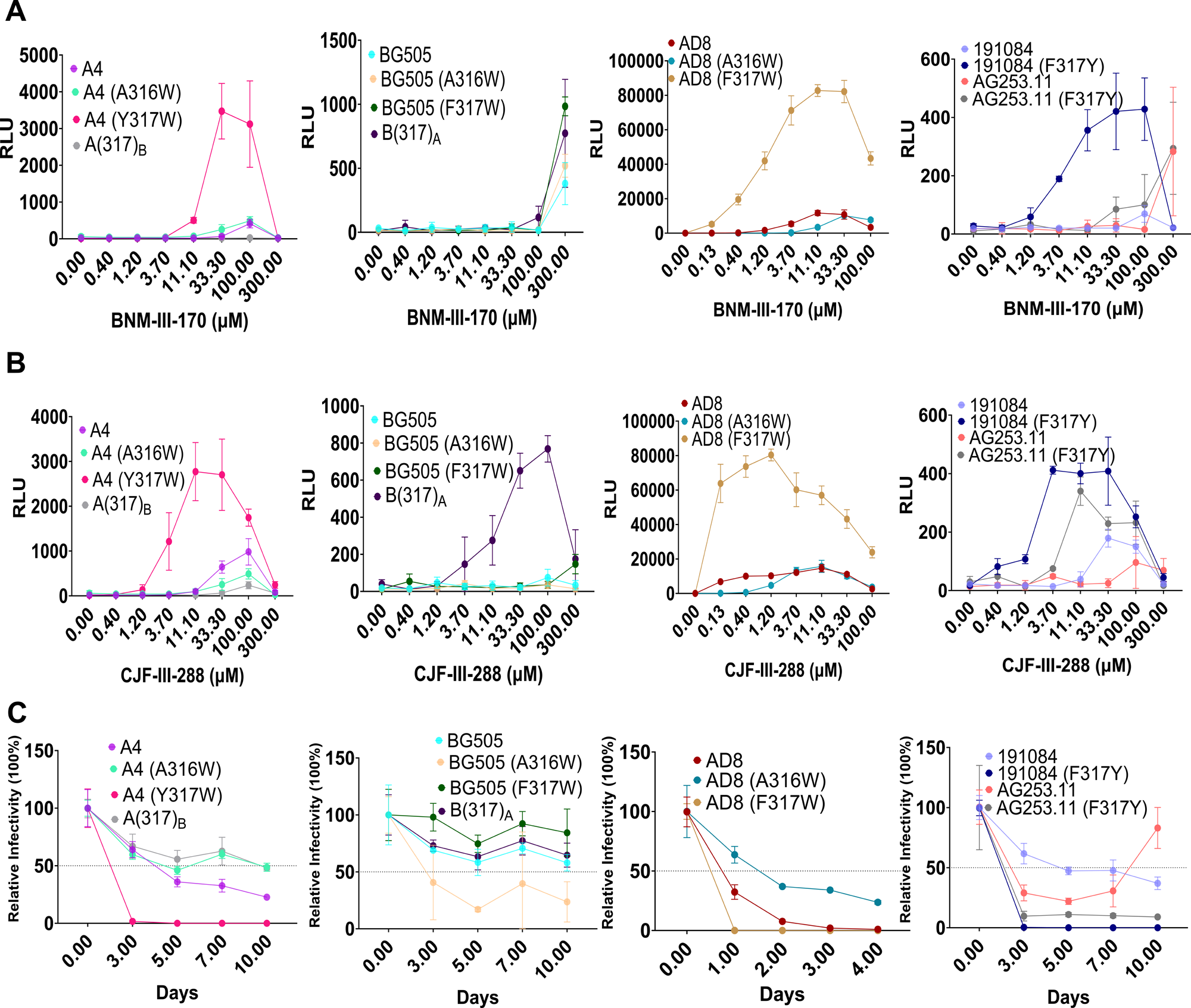
Activation of HIV-1 infection by CD4mcs and sensitivity to cold inactivation. Activation of infection of CD4-negative, CCR5-expressing cells by BNM-III-170 (A) and CJF-III-288 (B) was evaluated for HIV-1 variants with wild-type and mutant A4, BG505 and AD8 Envs. HEK 293T cells were transfected with plasmids expressing the indicated Envs, HIV-1 packaging proteins and a luciferase-expressing HIV-1 vector. After 48 hours, pseudoviruses were harvested and incubated with Cf2Th-CCR5 cells in 96-well plates. The plates were centrifuged at 600 × *g* for 30 min at 21°C. Medium containing serial dilutions of BNM-III-170 or CJF-III-288 was then added. After incubation at 37°C in a CO_2_ incubator for 48 hours, the cells were lysed and luciferase activity was measured. RLU, relative light units. (C) To evaluate cold sensitivity, pseudotyped viruses were produced as described in the Fig. 1A legend and were incubated on ice for the indicated times, after which the virus infectivity was measured. In A-C, the means and standard deviations of triplicate measurements are shown. The experiments were repeated with comparable results.

We evaluated the relationships among the viral phenotypes of the Env variants in this study. The sensitivities of the virus variants to inhibition by BNM-III-170 and CJF-III-288 were highly correlated (Fig. 6A); therefore, we used virus sensitivity to CJF-III-288 inhibition to evaluate potential relationships with other viral phenotypes. For viruses with V3 loop Env variants, there is a strong correlation between sensitivity to CJF-III-288 inhibition and activation by the CD4mcs (Fig. 6B).

**Fig 6:**
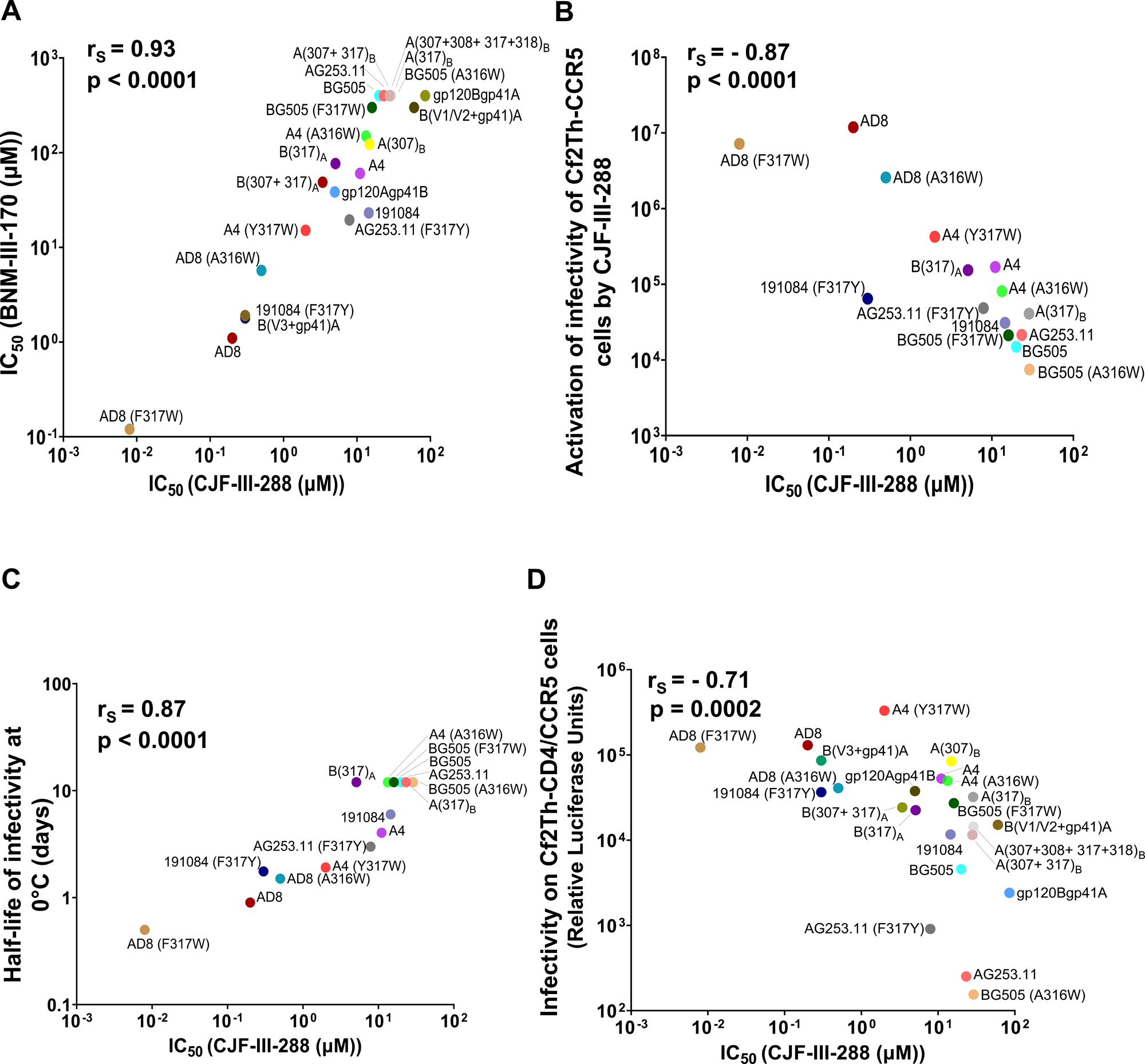
Correlations between viral phenotypes associated with Env variants. (A) Correlation between the IC_50_ values (in µM) for inhibition of Cf2Th-CD4/CCR5 cell infection by CJF-III-288 and BNM-III-170 for Env mutants in this study. (B) The relationship between inhibition of infection of Cf2Th-CD4/CCR5 cells (on the x axis) and activation of infection of Cf2Th-CCR5 cells (on the y axis) by CJF-III-288 is shown for the indicated pseudovirus variants. The x-axis values represent the IC_50_’s for each Env from Figs. 2A, 3A and 4A. The y-axis values represent the area of each peak calculated from the activation graphs in Fig. 5B. (C) The relationship between inhibition of infection of Cf2Th-CD4/CCR5 cells (on the x axis) by CJF-III-288 and sensitivity to cold exhibited by viruses with different Envs is shown. The x-axis values are as in B. The y-axis values represent the half-life of infectivity at 0°C obtained from the experiments shown in Fig. 5C. (D) The relationship between inhibition of infection of Cf2Th-CD4/CCR5 cells (on the x axis) by CJF-III-288 and infectivity of viruses with the different Envs is shown. The x-axis values are as in B. The y-axis values represent the viral infectivities from Figs. 1B, 2C, 3B and 4B. The Spearman rank-order correlation coefficients (r_S_) and *p* values are shown for each of the graphs.

### Cold sensitivity of mutant viruses

One of the phenotypes associated with alterations in the stability of the State-1 conformation of the HIV-1_AD8_ Env is a change in the sensitivity of the functional viral spike to inactivation by exposure to 0°C (63, 64, 78, 79). We evaluated the infectious half-life following incubation on ice for recombinant viruses pseudotyped by Env variants selected from the studies above. The infectivity of the viruses with the primary HIV-1 Envs exhibited a range of sensitivity to cold (Fig. 5C). Compared with the wild-type A4, AD8, 191084 and AG253.11 viruses, the viruses with Tyr or Trp substitutions at residue 317 were inactivated more rapidly at 0°C. By contrast, the A4 and AD8 viruses with the A316W change were more resistant to cold inactivation. For viruses pseudotyped with this panel of Env variants, there is a good correlation between sensitivity to CJF-III-288 inhibition and cold inactivation (Fig. 6C).

### Relationship between virus sensitivity to CD4mcs and infectivity

The relationship between the susceptibility to inhibition by CJF-III-288 and infectivity for the panel of HIV-1 Env variants used in this study is shown in Fig. 6D. Virus infectivity varied inversely with resistance to the CD4mc.

## DISCUSSION

Even when the highly resistant CRF01_AE recombinant HIV-1 strains are excluded, primary HIV-1 strains exhibit a 1000-fold range of IC_50_ values with respect to inhibition by CD4mcs (37, 57). The rank orders of sensitivities of HIV-1 Envs to different CD4mcs are highly correlated, indicating that Env sequences determine virus sensitivity to multiple CD4mcs (57). For the panel of Env variants used in this study, sensitivity to BNM-III-170 and CJF-III-288 correlated. A substantial fraction of the variation in primary HIV-1 sensitivity to CD4mcs is apparently determined by changes in Env elements outside of the known binding sites. Changes in Env triggerability, which is governed by the activation energy barrier between State 1 and downstream Env conformations, correlate with CD4mc sensitivity for carefully matched panels of HIV-1_AD8_ Env mutants (64). Additional studies are needed to evaluate the hypothesis that alterations in Env triggerability contribute to the variation in the sensitivity of natural HIV-1 strains to CD4mcs.

The difference in sensitivity of the A4 and BG505 Envs to CD4mcs was determined by the Phe/Tyr 317 polymorphism in the tip of the gp120 V3 loop. The tyrosine residue in the A4 Env is infrequent at this position and is less hydrophobic than the more common Phe, Leu or Trp residues. In available Env structures, the V3 loop projects towards the trimer axis at the membrane-distal apex (79–82) (Fig. 7A). Hydrophobic packing in the trimer apex has been suggested to contribute to the maintenance of the closed, pretriggered Env conformation (13, 29, 83). Based on this model, the more polar tyrosine 317 residue may disrupt packing of the trimer apex and predispose the pretriggered Env to open into downstream conformations. The phenotypes of the A316W and A317W mutants demonstrate that hydrophobic substitutions in this region may result in either increases or decreases in CD4mc resistance, respectively. If hydrophobic packing of the V3 tip is important for maintenance of a pretriggered state, the large tryptophan indole ring may be better accommodated at position 316 rather than 317. Trp 316 has been suggested to stabilize sgp140 SOSIP.664 Env trimers by stacking against Tyr 318 (77). In any case, Phe/Tyr 317 is located approximately 14 Angstroms from the nearest gp120 residue that contacts the CD4mcs and thus these V3 polymorphisms exert their phenotypic effects over a distance (Fig. 7B).

**Fig 7:**
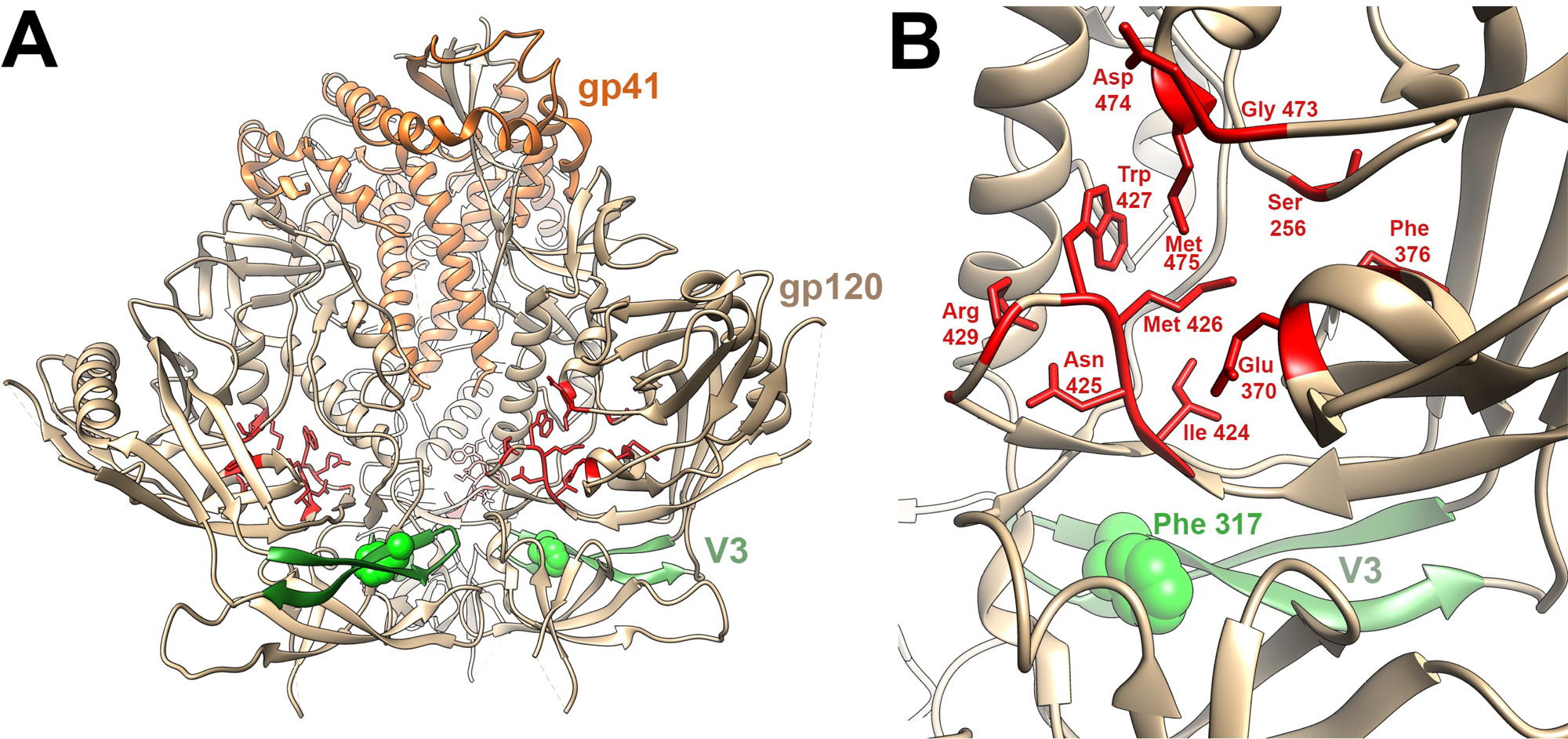
Relationship of the gp120 V3 loop and CD4mc binding site on an Env trimer structure. (A) A side view of the unliganded HIV-1_BG505_ sgp140 SOSIP.664 Env trimer (PDB 4ZMJ) (90) is shown, with gp41 (orange) at the top and gp120 (tan) at the bottom of the image. The gp120 V3 loop is colored green, with residue 317 shown in CPK representation. The gp120 residues within 3.5 Angstroms of the CD4mc CJF-III-288 in gp120 core co-crystals (PDB 8FM3) (57) are colored red and shown in stick representation. (B) Close-up view of the gp120 region encompassing the V3 loop (with Phe 317) and the CJF-III-288 binding site, represented as in A. The CJF-III-288 contact residues (in red) are completely conserved between the A4 and BG505 Envs, as is Ser 375, which is not visible from this perspective. In a single protomer of the HIV-1_BG505_ sgp140 SOSIP.664 Env trimer, Phe 317 is 13.5 Angstroms from the nearest residue (Ile 424) that contacts CJF-III-288. Phe 317 on the adjacent Env protomer is even further away from the CJF-III-288 binding site.

Our results reveal the functional consequences of naturally occurring polymorphisms in residue 317. In addition to conferring an increase in sensitivity to CD4mcs, Tyr 317 increased CD4mc-induced gp120 shedding and activation of infection of CD4-negative, CCR5-expressing cells. For the entire panel of Env mutants evaluated, virus sensitivity to CJF-III-288 inhibition correlated with CD4mc activation of infection, virus infectivity and cold sensitivity. The latter two Env phenotypes are evaluated in the absence of CD4mcs and therefore provide independent indicators of increased Env triggerability (26, 63, 64,78). An increase in triggerability specified by V3 residue 317 apparently contributes to the moderate increase in replicative ability and cold sensitivity of the A4 virus compared with the BG505 virus. The development of resistance to CD4mcs through down-modulation of Env triggerability is expected to incur a fitness cost reflected in the observed decreases in virus infectivity. It will be interesting to determine how properties associated with Env triggerability are related in a larger panel of primary HIV-1 strains.

In contrast to the changes in the gp120 Phe-43 cavity that arose during selection of a BNM-III-170-resistant virus (62), the alterations of the V3 loop observed here affected sensitivity to CD4mcs but not to BMS-806. Unlike CD4mcs, conformational blockers like BMS-806 and its analogues do not need to induce large-scale conformational changes in Env to exert their antiviral effect (12). As a result, BMS-806 analogues are less susceptible to Env alterations distant from the drug-binding site that lower triggerability. Thus, despite the proximity of the gp120 binding sites of CD4mcs and BMS-806 (84, 85), the distinct consequences of binding related to their antiviral mechanisms strongly impact potential HIV-1 pathways of resistance. An appreciation of the differences between these classes of gp120-directed virus entry inhibitors will guide potential applications.

## MATERIALS AND METHODS

### Plasmids

The *env* genes encoding the A4 and BG505 Envs and chimeric Envs were expressed using the pcDNA3.1 Env plasmid (Genbank accession numbers QPJ74671 for the A4 *env* and ABA61516 for the BG505 *env*). Additional primary HIV-1 Envs included in this study are from HIV-1_191084_ and HIV-1_AG253.11_ (Genbank accession numbers ADI62025 and ACC97453, respectively). For experiments in which the Envs were expressed in the context of an HIV-1 provirus, the *env* genes were cloned into the pNL4-3 proviral clone (86). Changes were introduced into the *env* genes using a Q5 site-directed mutagenesis kit (New England BioLabs). The presence of the desired mutations was confirmed by DNA sequencing.

### Cell lines and primary cells

293T cells were grown in Dulbecco’s Modified Eagle’s medium (DMEM) (Life Technologies, Wisent Inc.) supplemented with 10% fetal bovine serum (FBS) (Life Technologies, VWR) and 100 μg/ml of penicillin-streptomycin (Life Technologies, Wisent Inc.). Cf2Th-CD4/CCR5 cells stably expressing the human CD4 and CCR5 coreceptors for HIV-1 were grown in the same medium supplemented with 0.4 mg/ml of G418 and 0.2 mg/ml of hygromycin. Cf2Th-CD4/CCR5 cells stably expressing the CCR5 coreceptor of HIV-1 were grown in the same medium supplemented with 0.4 mg/ml of G418.

### Small-molecule HIV-1 entry inhibitors

The CD4-mimetic compounds (CD4mcs) BNM-III-170, CJF-III-192 and CJF-III-288 were synthesized as described previously (37, 57). The compounds were dissolved in dimethyl sulfoxide (DMSO) at a stock concentration of 10 mM and diluted to the appropriate concentration in cell culture medium for antiviral assays. BMS-806 was purchased from Selleckchem.

### Expression and processing of HIV-1 Env variants

HEK 293T cells were transfected transiently with plasmids encoding Envs, HIV-1 packaging proteins and a luciferase-expressing HIV-1 vector (26). Forty-eight hours later, the cell supernatant was cleared (600 × *g* for 10 min) followed by filtration through a 0.45-μm membrane. The viruses were pelleted by centrifugation at 14,000 x *g* for 1.5 hours at 4°C and lysed. The cell and virus lysates were Western blotted and probed with a goat polyclonal anti-gp120 antibody (Invitrogen), the 4E10 anti-gp41 antibody (NIH HIV Reagent Program) or a rabbit anti-Gag antibody (Abcam). The cell lysates were also Western blotted and probed with a rabbit anti-hsp70 antibody (Santa Cruz).

### Production of recombinant pseudoviruses expressing luciferase

As described previously (26), 293T cells were transfected with pSVIIIenv plasmids expressing Env variants, the pCMVΔP1Δenv HIV-1 Gag-Pol packaging construct and the firefly luciferase-expressing HIV-1 vector at a 1:1:3 µg DNA ratio using effectene transfection reagent (Qiagen). Recombinant, luciferase-expressing viruses capable of a single round of replication were released into the cell medium and were harvested 48 h later. The virus-containing supernatants were clarified by low-speed centrifugation (600 × *g* for 10 min) and used for single-round infections.

### Virus infectivity, inhibition, and cold sensitivity

Single-round virus infection assays were used to measure the ability of the Env variants to support virus entry, as described previously (26). To measure the infectivity of the Env pseudotypes, recombinant viruses were added to Cf2Th-CD4/CCR5 target cells expressing CD4 and CCR5. Forty-eight hours later, the target cells were lysed and the luciferase activity was measured.

To measure virus inhibition, the compounds to be tested were incubated with pseudoviruses for 1 hour at 37°C. The mixture was then added to Cf2Th-CD4/CCR5 target cells expressing CD4 and CCR5. Forty-eight hours later, the target cells were lysed and the luciferase activity was measured.

To evaluate the cold sensitivity of the Env variants, pseudotyped recombinant viruses were incubated on ice for various lengths of time prior to measuring their infectivity, as described previously (26, 27, 63, 64, 78).

### gp120 shedding

To measure CD4mc-induced gp120 shedding, 293T cells were transfected transiently with pNL4-3 provirus constructs expressing the A4, BG505, A(317)_B_ and B(317)_A_ Envs. The transfected cell supernatants were harvested 48 hours later, clarified by low-speed centrifugation (600 × *g* for 10 min) and filtered through a 0.45-μm membrane. The viruses were pelleted by centrifugation at 14,000 x *g* for 1.5 hour at 4°C and resuspended in 100 µl of 1X PBS. The viruses were incubated with different concentrations of CJF-III-288 for 2.5 hours at 37°C, followed by centrifugation at 14,000 x *g* for 1.5 hours at 4°C. The virus pellets were lysed in 1X LDS buffer. The supernatants containing shed gp120 were bound to Galanthus nivalis lectin (GNL) beads (Thermo Fisher Scientific). The glycoproteins captured on the beads and the lysates of virus pellets were Western blotted and probed with a goat anti-gp120 antibody, the 4E10 anti-gp41 antibody or a rabbit anti-Gag antibody. The Western blots were quantified using Image J software (87).

### Activation of virus infection by CD4mcs

Pseudoviruses were incubated with CD4-negative, CCR5-expressing Cf2Th-CCR5 cells in 96-well plates. The plates were centrifuged at 600 × *g* for 30 min at 21°C. Medium containing serial dilutions of CD4mc was then added. Forty-eight hours later, cells were lysed, and luciferase activity was measured.

### Statistics

The concentrations of HIV-1 entry inhibitors that inhibit 50% of infection (IC_50_ values) were determined by fitting the data in five-parameter dose-response curves using GraphPad Prism 8. Spearman rank-order correlation coefficients (r_S_) and p values were calculated using VassarStats (88).

## ACKNOWLEDGMENTS

We thank Ms. Elizabeth Carpelan for manuscript preparation. Antibodies against HIV-1 were kindly supplied by Dennis Burton (Scripps), Peter Kwong and John Mascola (Vaccine Research Center NIH), Barton Haynes (Duke University), Hermann Katinger (Polymun), James Robinson (Tulane University), and Marshall Posner (Mount Sinai Medical Center). We thank the NIH HIV Reagent Program for providing reagents.

This work was supported by grants from the National Institutes of Health (grants AI145547, AI124982, AI129017, AI164562, AI150471, AI148379, AI150322, AI129769 and AI176904), by an HIV Cure Research Grant from Gilead Sciences, and by a gift from the late William F. McCarty-Cooper.

